# Engineered Fluorescent Carbon Dots for Selective Cellular Bioeffects: A Comparative Study of Cancer and Normal Cells

**DOI:** 10.1101/2025.06.02.657379

**Authors:** Ankesh Kumar, Pankaj Yadav, Dhiraj Bhatia

## Abstract

Cancer remains one of the most critical global health challenges, with early detection being essential for effective treatment and improved survival rates. However, conventional diagnostic tools often fail to detect cancer at early stages due to limitations such as low sensitivity, high cost, and dependence on large tumour presence. Commercial dyes used for imaging are frequently hindered by poor water solubility, toxicity, instability, and high expense. In recent years, CDs have emerged as promising fluorescent probes due to their nanoscale size, tenable surface properties, strong fluorescence, and excellent biocompatibility. make them applicable to various biological applications such as bioimaging, drug delivery, and tissue engineering. In this study, CDs were synthesized using citric acid and ascorbic acid as carbon sources via a reflux method at 130 °C for 12 hours in a water-ethanol medium. The resulting CDs exhibited high water solubility, strong photostability, and low toxicity. Notably, they effectively distinguished cancerous cells from normal cells. And showing higher uptake in cancer cells due to increased membrane permeability and metabolic activity. Higher uptake means more accumulation within cells that leads to an increment in fluorescence intensity, based on the fluorescent intensity distinguishing the cancer cells and normal cells. These findings highlight the potential of CDs as cost-effective, biocompatible imaging agents for cancer diagnosis and cellular studies.

## 1. INTRODUCTION

Cancer is the leading cause of death worldwide. Early detection of the disease significantly enhances treatment success, increases survival rates, reduces complications, and lowers healthcare costs[1]. Cancer ranks as the second leading cause of death worldwide, resulting in approximately 9.6 million fatalities, which translates to 1 in every 6 deaths, in 2018[2–4]. Among men, the most prevalent types of cancer include lung, prostate, colorectal, stomach, and liver cancer, whereas breast, colorectal, lung, cervical, and thyroid cancers are the most common for women[5].

Cancer treatment has a higher success rate when the disease is detected early[1]. Early identification leads to improved survival rates, reduced morbidity, and lower treatment costs. For early cancer detection, Various dyes are available for cancer detection[6], including fluorescein sodium[7], methyl blue[8], and Indocyanine green (ICG)[9]. However, these dyes come with their own set of limitations, such as low fluorescence, poor water solubility, cytotoxicity, instability, complicated synthesis processes, and high costs[10].In addition, Antybody-based fluorophores are used for early cancer detection. One notable example is the Anti-CEA Antibody[11], which effectively stains gastric and colorectal tissues by targeting the carcinoembryonic antigen[12] (CEA) present in cancer cells. Another example is the NIR fluorescence dye (BM-105), which helps to localize tumours, offering essential guidance during surgical procedures. PSMA-targeted probes[13] represent an important category, specifically designed to target prostate cancer by binding to the prostate-specific membrane antigen (PSMA) found on cancer cells. These probes are valuable for both imaging and radiopharmaceutical therapy. However, antibody-based fluorophores do face challenges, including immunogenicity, delays in antigen identification, photobleaching, and high costs[14–16]. Enzyme receptor-based fluorophores also show promise for cancer detection. For instance, MMP-Activatable[13] Probes are used for breast cancer and are activated by matrix metalloproteinases (MMPs) that cleave peptide linkers, leading to a loss of FRET and subsequent fluorescence activation. Similarly, Cathepsin-Activatable Probes[17] target Cathepsin D, which cleaves a peptide substrate to release the BODIPY fluorophore, enabling pH-insensitive detection in lysosomes and phagosomes. Despite their potential, enzyme-based fluorophores are not without limitations, such as structural instability, complex synthesis, high cost and limited tissue penetration[18].

Early detection of cancer can significantly reduce the death rate. However, this poses difficulties since the tumour needs to reach a detectable size. At present, cancer detection methods primarily rely on imaging techniques[19] such as computed tomography (CT)[20], X-ray[21], magnetic resonance imaging (MRI)[22], and ultrasound[23], which encounter challenges including high radiation exposure, low signal-to-noise ratios, and high costs. Therefore, there is a pressing need for the creation of more effective and novel approaches for the early detection of cancer[24]. Fluorescent nanoparticles offer a more effective means of selectively targeting cancerous cells. The advancement of novel nanoparticle-based fluorophores should include outstanding characteristics such as high photostability, biocompatibility, easy synthesis, and excellent fluorescence to differentiate between cancerous and normal cells[25].

In recent years, fluorescence nanoparticles[26,27] have gained significant attention due to their unique property. Among them, Carbon dots have excellent optical characteristics, such as biocompatibility, excellent photostability, strong fluorescence, easy synthesis, and cost-effectiveness[28–33].The cross-linking of DNA strands to form DNA nanocages[34–38]. These nanocages are used in drug delivery and cell transplantation. these DNA-based cargoes[39,40] have unique properties such as large surface area, high stability, high drug loading capacity, and enhanced ability for biological interactions[41–44]. They are more prominent for controlled and targeted drug delivery drug delivery[45–48].

There is growing excitement around using DNA-based carbon quantum dots (CQDs) for bioimaging owing to their great biocompatibility, flexible ability, excellent fluorescence, surface modifications[49–51]. Usually obtained by hydrothermal or bottom-up synthesis using DNA or similar molecules, these nanoparticles fluorescence strongly, are water-soluble, have low toxicity and work well for imaging in cells and the body[52,53]. Because of their structure and functional surface, core-shell nanoparticles can easily attach ligands, antibodies or medicines, making it easier to target and see particular structures inside cells[54–58]. DNA-based CQDs can enter cells well, collect inside tumours through a peculiar property of the tumour and supply fluorescent, real-time signals for watching cellular activities or how diseases evolve[55,59]. In contrast to traditional quantum dots made from metals or organic dyes, DNA-based CQDs are very photostable[60], which helps with long-term imaging and clinical diagnostics[61,62]. Furthermore, if these dots are doped with heteroatoms or combined with polymers, this can lead to enhanced optical properties and better targeting, permitting them to do more in advanced biomedical imaging and theragnostic uses[63–65]. DNA-based carbon quantum dots shine as a flexible, safe and reasonably priced base for creating new bioimaging tools[66]. in addition, research shows here for the first time the production of fluorescent DNA hydrogel using just a slight number of C-Dots as the crosslinker[67]. Special attention was given to the optimization of the dots so they would selectively detect dopamine using electrochemistry. The many benefits allow DNA nanostructures to be designed for medical use in drug delivery, treatment, imaging and detection[68–78]. Nanotechnology and bioanalysis can be enhanced by modified functional DNAs which appear useful in assembling DNA nano-architectures for use in biotechnology and medicine[79,80].

Carbon dots are emerging nanoparticles with a size range of 2-10nm and the shape and morphology of quasi-spherical[81]. Carbon Dots became even more interesting to scientists once it was seen that they have excellent fluorescence[82], noticed during the first purification of single-walled carbon nanotubes in 2004[83,84]. Researchers have identified different ways fluorescence occurs in CDs such as in the molecular state, because of quantum confinement, due to surface defects and due to enhanced emission after cross-linking[85]. The reason they are becoming[86] popular in biomedicine is that they are bright, fluorescent, and biocompatible, making them useful for cancer therapy, gene delivery, imaging, sensing and drug delivery. For nanomaterials and nanostructures used in bioimaging, they must be taken up by both cells and tissues. Endocytosis is the route that lets any nanomaterial enter the cells[87–96]. As a result, one must understand the cellular systems that help take in CDs. The body uses mainly phagocytosis and pinocytosis to take in macromolecules and nutrients from outside the cells. While the former engulfs bigger particles using phagocytosis, the latter is involved in taking up smaller ones that are not engulfed by the larger cells[97].

Earlier researchers used modified CDs to distinguish between normal cells and cancer cells; this study explores the unmodified or unfunctionalized CDs that were used for distinguishing Between normal cells and cancer cells. We address CDs, more uptake shows in cancer cells as compared to normal cells. That led to the increment in fluorescent intensity. based on the fluorescent intensity distinguish between normal cells and cancer cells. There is a key factor that influences higher uptake in cancer cells, Cancer cells are located within tumour tissues that exhibit leaky blood vessels and inadequate lymphatic drainage[98], which facilitates a higher accumulation of core-shell carbon dots in these tumours compared to normal tissues through a passive targeting mechanism[99] that is less effective in healthy cells. Additionally, the negatively charged membranes[100] of cancer cells, due to the presence of excess phosphatidylserine, can enhance the interaction with CDs designed with zwitterionic or pH-responsive surfaces[101] that may switch to a positive charge in the acidic environment of tumors, resulting in greater uptake. Moreover, cancer cells tend to overexpress certain transporters, such as LAT1 (L-type amino acid transporter 1)[102], allowing for the active internalization of CDs that are functionalized with amino acid mimics, a feature that normal cells do not possess. The metabolic landscape also differs, as elevated levels of glutathione (GSH) in cancer cells improve the fluorescence signal of CDs/Fe^3+^ complexes, facilitating more effective tumor imaging, while normal cells display lower GSH levels and weaker signals. Furthermore, the rapid proliferation of cancer cells leads to increased endocytic activity, which significantly boosts CD internalization; for instance, CdSe@CdS nanocrystals exhibit four times higher fluorescence intensity in HeLa cancer cells compared to normal cells[103].

**Figure 1.**
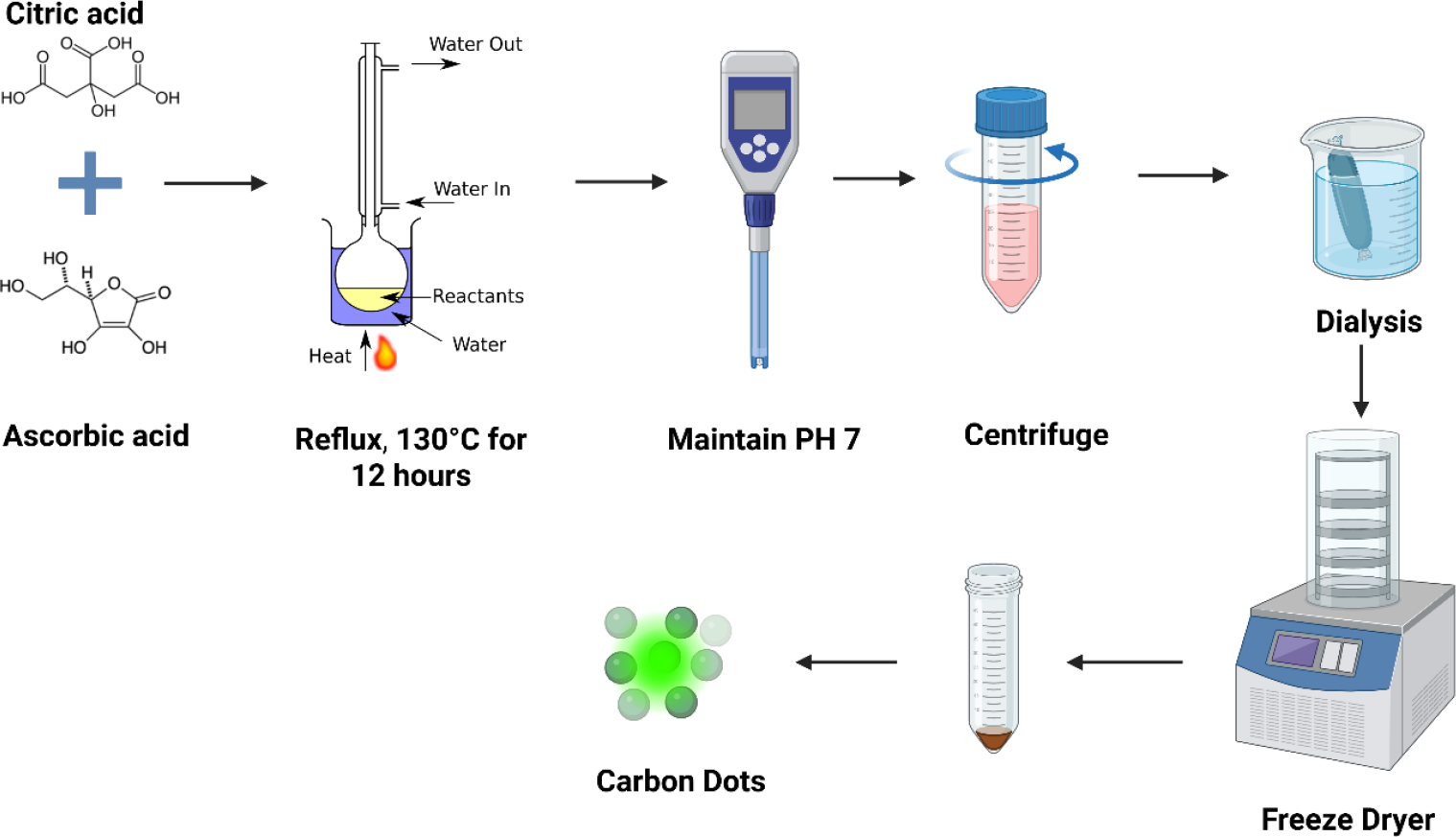
Schematic representation and synthesis of carbon dots differentiate normal cells from cancer cells.

## 2. MATERIALS AND METHODS

### 2.1. Materials

L-Ascorbic acid (99.5%) from Loba chemicals Citric acid anhydrous (extra pure) was purchased from SRL chemicals, ethanol (>99.9%), Silicon oil was purchased from X-chemicals was supplied by Changshu Hongsheng Fine Chemicals Co., Ltd., 0.22-micrometer filter, 3.5 KDa snakeskin membrane thermos fisher, Milli-Q water from Merck Millipore. Cell culture dishes and dimethyl sulfoxide (DMSO) were obtained from HiMedia. Dulbecco’s Modified Eagle Medium (DMEM), 0.25% trypsin-EDTA, and fetal bovine serum (FBS) were purchased from Gibco. All reagents used were of analytical grade and required no further purification. MDA-MB-231 (Breast Cancer), HeLa (Cervical Cancer), Rpe1 (Retinal pigment epithelial cells), HEK293T (Human Embryonic Kidney cells, NIT3T3 (mouse NIH/Swiss embryo) were maintained in DMEM, the cells were kept at a temperature of 37 °C with 5% CO2 in a humidified incubator.

### 2.2. Methods

#### 2.2.1. Synthesis of Carbon Dots

Dissolve 1 g each of ascorbic acid and citric acid in twice the volume of water and an equal volume of ethanol. The mixture should then be heated at 130ºC for 12 hours. After cooling overnight. Collect the resulting solution in a 50 mL Falcon tube and adjust the pH to 7 using 10 M NaOH. Then centrifuge the mixture for 10 minutes at 10,000 rpm at room temperature. CDs obtained were dialyzed against deionized water using a 3.5 kDa snakeskin membrane. Finally, the fluorescent carbon dots were collected into a 50 mL Falcon tube after the solvent was removed through lyophilization.

#### 2.2.2. Characterization of Green fluorescent carbon dots

The optical properties of the CDs, including UV-Vis absorbance and fluorescence emission, were measured using a Spectrocord-210 Plus from Analytokjena (Germany) and an FP-8300 Jasco spectrophotometer (Japan), respectively. For the stock solution, 200 µL of a 1 mg/mL sample was dissolved in 1 mL of Milli-Q water in a 1 mL tube. After preparing the solution, 1 mL of the diluted sample was transferred into a cuvette for analysis. Following the analysis, the data obtained was collected and plotted using Origin software for further interpretation.

For additional characterization, the CDs at pH 7 were lyophilized to obtain a powder. The FTIR spectra were then recorded in the range of 400 cm-1 to 4000 cm-1 using a PerkinElmer Spectrum 2 in ATR mode.

The morphology-based characterization was carried out using Atomic Force Microscopy (AFM). To prepare the sample, a 1cm^2^ mica sheet was adhered to a glass slide using nail polish and then cleaved using cello tape. The sample was initially prepared at a concentration of 1 mg/ml in milli-Q water, followed by a dilution of 20 µl in 1 ml of milli-Q water. Subsequently, 5 to 7 µl of the diluted sample was drop-cast onto the mica sheet. After allowing the sample to settle, the excess was removed, and the surface was washed twice with ultrapure water. The samples were dried thereafter inside a vacuum-desiccator to remove any water content. Finally, the samples were imaged using the BIO AFM (Bruker’s Nanowizard Sense plus Bio-AFM) in tapping mode under ambient conditions. For the imaging process, a sharp silicon cantilever ScanAsyst-Air from Bruker was employed.

#### 2.2.3. Cell culture and cellular uptake assay

RPE1 and MDA-MB231 cells were plated on a 10 mm cover slide in 4-well plates, with 3 × 10^5 cells added to each well in a growth medium (DMEM with 10% FBS, 1x antibiotic mix (Penstrep), HEPES, Sodium Bicarbonate, and Sodium Pyruvate). The cells were incubated for 24 hours in a 5% CO2 atmosphere at 37ºC before the experiment. After removing the complete medium, the cells were washed three times with 1× PBS buffer and then incubated in serum-free medium for 15 minutes at 37 °C in a 5% CO2 humidified incubator. Following the washes, the cells were treated with CDs at concentrations of 0, 50, 100, and 200 µg/ml. The treated cells were then fixed for 15 minutes at 37 °C using 4% paraformaldehyde and rinsed three times with 1× PBS. Subsequently, the cells were permeabilized with 0.1% Triton-X100 and stained with 0.1% phalloidin to visualize the actin filaments. After three additional washes with 1× PBS, the cells were mounted on slides with Mowiol and DAPI to label the nuclei.

#### 2.2.4. MTT assay

To evaluate the cytotoxic effects of CDs, approximately 1×10^4^ RPE1 (Retinal Pigment Epithelial) and MDAMB cells were seeded in a 96-well plate with 100 µl of media consisting of DMEM, 10% FBS, 1x antibiotic (Penstrep), HEPES, Sodium Bicarbonate, and Sodium Pyruvate. The cells were allowed to acclimate for 24 hours in a 5% CO2 atmosphere at 37ºC before performing the experiment. The following day, the cells were washed with PBS and treated with increasing concentrations of CDs (10, 20, 30, 40, 50, 60, 70, and 80 µg/mL) for 24 hours in a serum-free medium. After treatment, the culture medium was removed, and 100 µl of DMEM containing MTT (5 mg/mL) was added, followed by incubation for 3 to 4 hours to assess the mitochondrial dehydrogenase activity of live cells. The medium was then aspirated again, and 100 µl of DMSO was added to each well to dissolve the purple formazan crystals. Absorbance was measured at 565 nm using a Byonoy’s Absorbance 96 microplate reader. The experiment was performed in triplicate, with results normalized to the corresponding wells containing DMSO, while the wells without treatment served as controls for calculating the percentage of cell viability for each well. The untreated CDs served as the control to determine the % cell viability of each well. The cell viability percentage was calculated using the following formula:

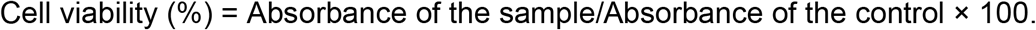

#### 2.2.5. Confocal imaging

Confocal imaging of fixed cells was conducted using a Leica TCS SP8 confocal laser scanning microscope (CLSM, Leica Microsystem, Germany) with a 63x oil immersion objective. Various fluorophores were excited using specific lasers: Hoechst at 405 nm, CDs at 488 nm, and phalloidin at 647 nm. Image quantification was carried out using Fiji software. For the analysis, whole cell fluorescence intensity was assessed at maximum intensity projection, with background subtraction applied, and the intensity values were normalized against those of unlabelled cells. A total of 32 to 40 cells were analyzed from the acquired z-stacks for each experimental condition.

#### 2.2.6. Statistical analysis

Statistical evaluation was conducted utilizing GraphPad Prism software (version 8.0.2). All data are presented as means ± standard deviation (SD) or means ± standard error derived from two independent experiments. p-values were determined through one-way ANOVA and two-tailed unpaired Student’s t-tests, with a confidence interval of 95%.

#### 2.2.7 Image Processing and Statistical Analysis

Image analysis was performed using Fiji (ImageJ). Z-stack images were transformed into two-dimensional formats utilizing the Z-projection function, specifically applying the maximum intensity projection technique. Background correction was executed by subtracting the mean background intensity from each image. Subsequently, quantitative assessments of integrated density and raw integrated density were collected.

## 3. RESULTS AND DISCUSSION

### 3.1. Characterization of CDs

Citric acid and ascorbic acid were dissolved in a water-ethanol mixture (2:1 ratio) and subjected to reflux heating at 130 °C for 12 hours. After cooling the reaction mixture to room temperature, the pH was adjusted to 7 using 10 M NaOH. The solution was then centrifuged to remove any insoluble residues, and the supernatant was collected. This was followed by dialysis to remove small molecular impurities. Finally, the purified solution was lyophilized to obtain dry carbon dots powder for further characterization and cell culture studies. illustrated in figure1.

**Figure 1.**
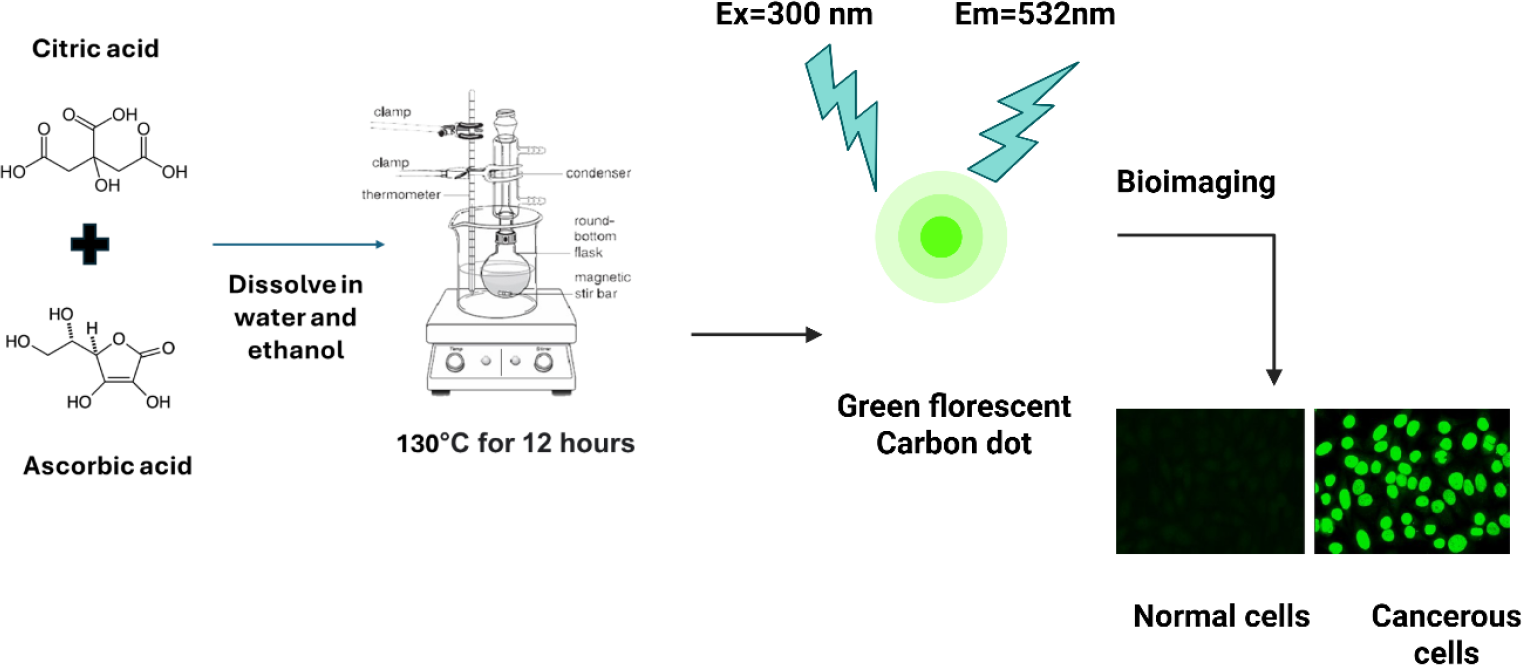
Synthesis of fluorescent CDs. Dissolve equal amounts of ascorbic acid and citric acid in twice their volume of water, along with an equal volume of ethanol. Heat the mixture at 130ºC for 12 hours. After cooling overnight, transfer the resulting solution into a 50 mL Falcon tube and adjust the pH to 7 using 10 M NaOH. Centrifuge the mixture at room temperature for 10 minutes at 10,000 rpm. The carbon dots obtained are then dialyzed against deionized water using a 3.5 kDa snakeskin membrane. Finally, collect the fluorescent carbon dots into a 50 mL Falcon tube after removing the solvent through lyophilization.

The optical properties of the generated Carbon dots were examined through the analysis of UV spectra and fluorescence spectra. In UV absorbance spectra, observe a 265 nm peak. The 265 nm corresponds to the carbonic core centre. The band around 265 nm corresponds to the n−π* transition C═O bond, as shown in Figure 2 (a). The photoluminescence spectra excitation 300nm revealed that maximum emission intensity was observed at a wavelength of 532 nm when excited at 450 nm wavelength, confirming the green fluorescence as shown in Figure (b). To confirm their fluorescence properties, the GCDs were placed in a UV light chamber. Under daylight, no visible fluorescence was observed, but under UV light, the GCDs emitted a bright green fluorescence. This fluorescence arises from the electronic transitions within the carbon dots, as shown in Figure 2(c). furthermore, Dynamic Light Scattering analysis shows the hydrodynamic size distribution of GCDs, with a dominant peak at ∼3.9 nm indicating the average particle size, as shown in Figure 2(d). The FTIR spectrum of GCDs displays several characteristic absorption bands that indicate the presence of various functional groups. A broad peak around 2929 cm−^1^ corresponds to the O–H stretching vibration of carboxylic acids. The peak at 1564 cm−^1^ is attributed to C=C stretching in aromatic rings and possibly N–H bending, suggesting the presence of aromatic and amine groups. Additionally, peaks at 1385 cm−^1^ and 1079 cm−^1^ are associated with C–O stretching vibrations from alkoxy and phenol or acyl groups, respectively. The presence of alkenes and aromatic structures is further confirmed by bending vibrations at 911 cm−^1^ and 839 cm−^1^, corresponding to alkene sp^2^ C–H and aromatic sp^2^ C–H bending. These features indicate that the GCDs contain carboxylic, aromatic, phenolic, alkene, and possibly amine functionalities, typical of functionalized carbon-based nanomaterials, as shown in Figure 2(e). AFM images Figures (f) to (i) displayed the surface morphology and distribution of GCDs with scale bars of 50 nm and. These images reveal that the GCDs are uniformly dispersed and exhibit a nearly spherical shape with smooth surface features. The mean size length, height, and width are 7.61,8.45,7.72nm, respectively. that confirms that the CD size is>10nm and the shape is spherical.

**Figure 2.**
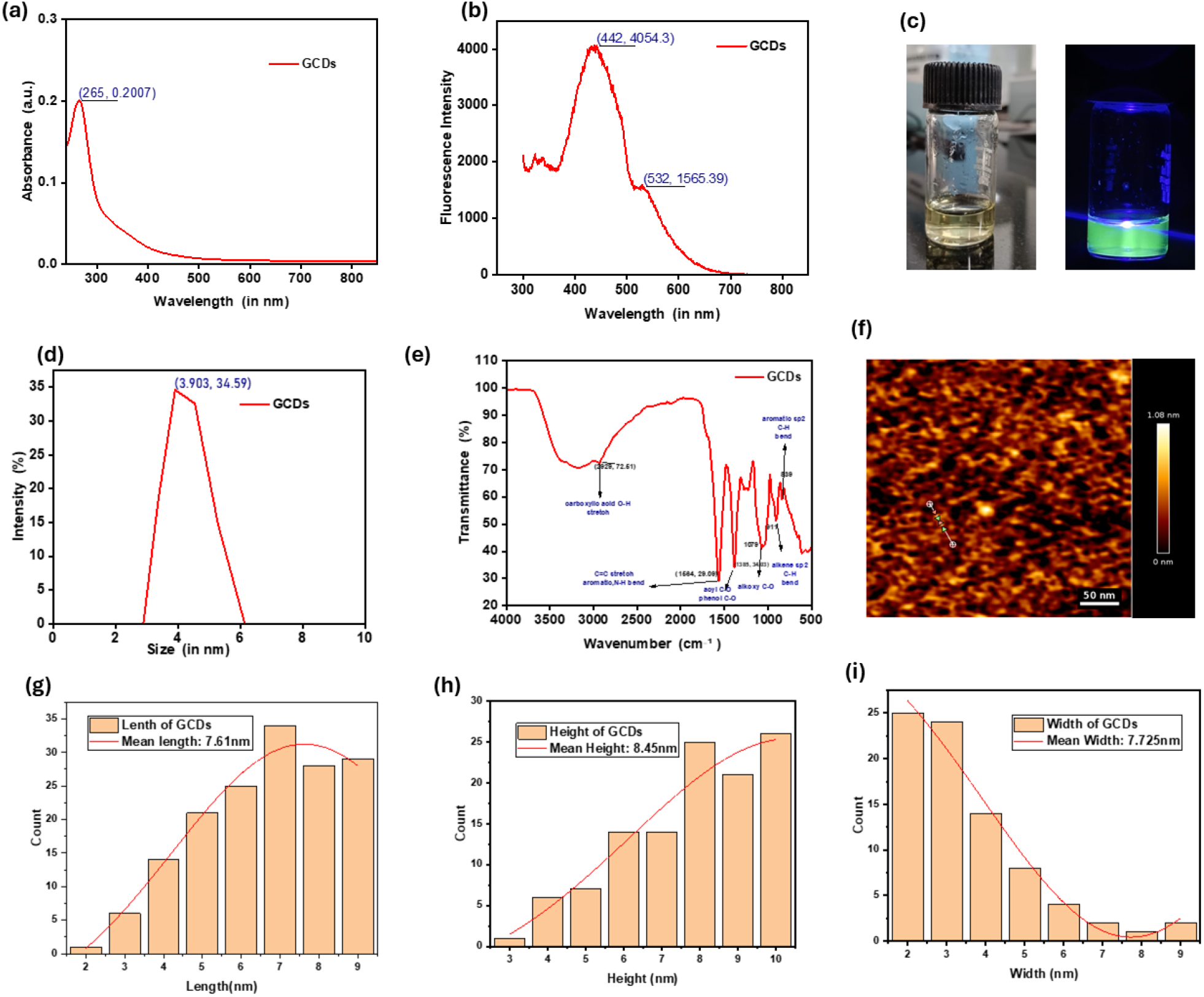
Characterization of CDs: figure 2 (a) The absorption peak observed at 265 nm is linked to the carbon core of the GCDs, the bands near 265 nm are related to the n→π* transitions of C=O bonds. Figure 2(b) The emission spectrum of GCDs excited at 300 nm shows prominent peaks at 442 nm (maximum intensity ≈ 4054 a.u.) and a shoulder at 532 nm. The observed fluorescence peaks are attributed to π–π* transitions within the sp^2^ domains of the carbon core and n–π* transitions associated with surface functional groups. Figure 2((c) The fluorescence of GCDs in room light (on the left) and under UV light (on the right). Under daylight, no visible fluorescence was observed, but under UV light, the GCDs emitted a bright green fluorescence. This fluorescence arises from the electronic transitions within the carbon dots. Figure (d) Dynamic Light Scattering analysis showing the hydrodynamic size distribution of GCDs, with a dominant peak at ∼3.9 nm indicating the average particle size. Figure (e) The FTIR spectrum of GCDs displays several characteristic absorption bands that indicate the presence of various functional groups. A broad peak around 2929 cm−^1^ corresponds to the O–H stretching vibration of carboxylic acids. The peak at 1564 cm−^1^ is attributed to C=C stretching in aromatic rings and possibly N–H bending, suggesting the presence of aromatic and amine groups. Additionally, peaks at 1385 cm−^1^ and 1079 cm−^1^ are associated with C–O stretching vibrations from alkoxy and phenol or acyl groups, respectively. The presence of alkenes and aromatic structures is further confirmed by bending vibrations at 911 cm−^1^ and 839 cm−^1^, corresponding to alkene sp^2^ C–H and aromatic sp^2^ C–H bending. AFM images Figures (f) to (i) displayed the surface morphology and distribution of GCDs with scale bars of 50 nm, The mean size length, height, and width are 7.61,8.45,7.72nm, respectively. that confirms that the CD size is>10nm and the shape is spherical, N (no of Carbon Dots) 75.

#### Zeta potential

The surface charge of the GCDs was analyzed using dynamic light scattering (DLS) by measuring the zeta potential at various pH levels: 2.74, 3.59, 7.0, and 10.3. The results indicated that as the pH increased, the zeta potential of the GCDs decreased, which reflects a reduction in surface charge. This trend suggests that at higher pH levels, the surface functional groups on the GCDs become less protonated. Consequently, this leads to a decrease in positive surface charge or an increase in negative charge, depending on the specific nature of the surface chemistry.

**Table No 1.** The zeta potential of GCDs at different pH values. The surface charge of the GCDs was analyzed using dynamic light scattering (DLS) by measuring the zeta potential at various pH levels: 2.74, 3.59, 7.0, and 10.3. The results indicated that as the pH increased, the zeta potential of the GCDs decreased, which reflects a reduction in surface charge.

| PH | 2.74 | 3.59 | 7 | 10.3 |
| --- | --- | --- | --- | --- |
| Zeta potential (mV) | +ve 8.943 | + ve 4.69 | + ve 2.87 | -ve22.7 |

### 3.2. Cytotoxic study of CDs

Since the knowledge about the cytotoxic effect caused by the system in normal (non-cancerous) cells and non-cancer cells is crucial, the cytotoxicity was studied using an MTT assay. The experiment was performed in both Cancerous (MDAMB231) and non-cancerous (RPE1) cell lines.

The cytotoxicity and biocompatibility of GCDs were examined using standard cell lines, specifically the retinal epithelial cell line RPE1 and a breast cancer cell line MDAMB231. In Fig. (a) RPE1 cells maintained a high viability of over 80% across all tested concentrations of GCDs (ranging from 0 to 100 µg/mL), demonstrating excellent biocompatibility and low cytotoxicity towards normal cells. In contrast, Fig. (b) illustrates the viability of the breast cancer cells MDAMB231 significantly decreased in a dose-dependent manner, dropping to nearly 50% at higher concentrations (up to 80 µg/mL). These results indicate that GCDs possess selective cytotoxicity; they are non-toxic to normal cells while exhibiting potent anticancer activity against cancer cells, highlighting their potential as biocompatible and effective agents for cancer therapy.as shown in figure 3.

**Figure 3.**
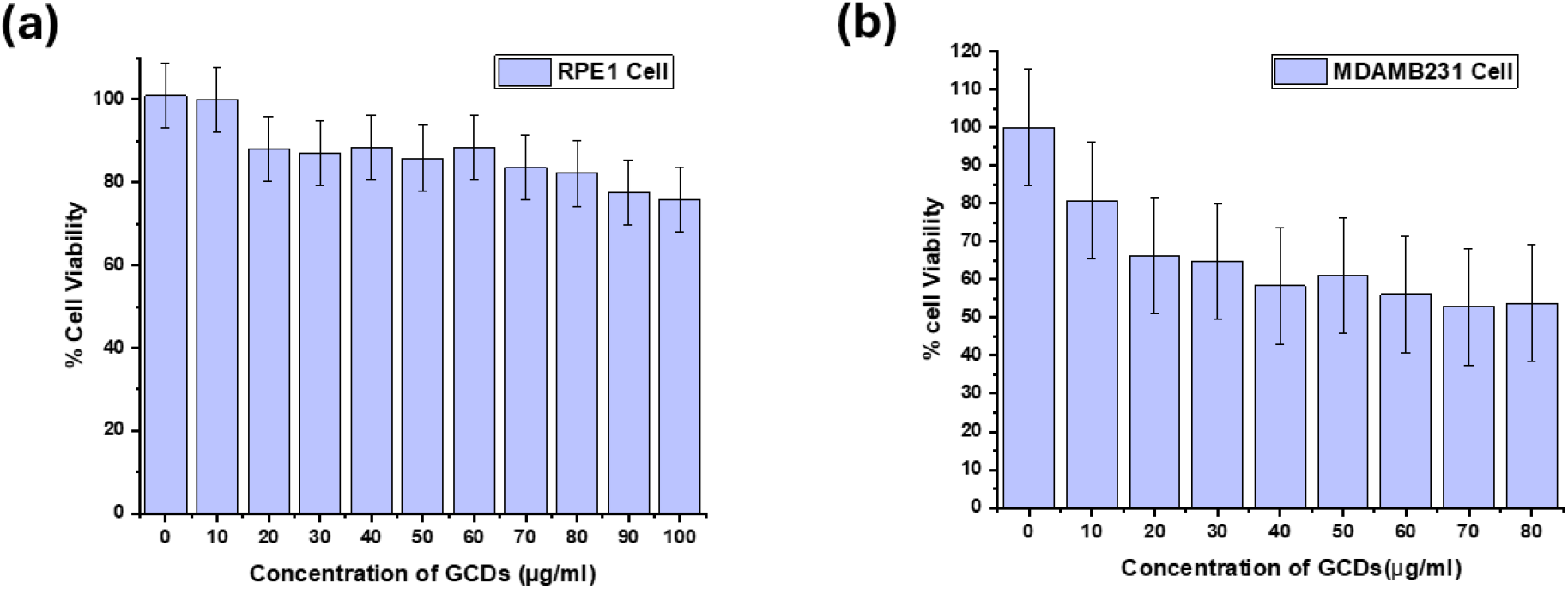
Cytotoxicity of GCDs in mice RPE1Cell and MDAMB231 cells. (a) Normal cells (b) Cancerous cells.

### 3.3. Cellular uptake of CDs

Literature reports indicate that GCDs can internalize into cells within 20 minutes. Based on this, we hypothesize that the uptake of GCDs is higher in cancer cells compared to normal cells. Cellular uptake was assessed using confocal microscopy. Fluorescence images were analysed using Image J software to quantify fluorescence intensity, which served as a proxy for GCD internalization. The quantified fluorescence data were plotted and statistically analysed using GraphPad Prism. As per the previous report, the GCDs have anti-cancerous properties. The fluorescence GCDs, along with marker DAPI (stain nucleus) and Phalloidin A647 (stain cytoskeleton of the cells). Aided to visualize cellular morphology and physiology.

Confocal images illustrate the concentration-dependent internalization of GCDs in RPE1 cells, with a scale bar of 20 µm. The fluorescence intensity of the CDs in RPE1 cells was quantified. In the case of normal cells (RPE1), there was an increased cellular uptake at concentrations of 100 and 200 μg/ml of Gcds compared to control and 50 μg/ml. The cellular uptake study is shown in the figure 4 above.

**Figure 4.**
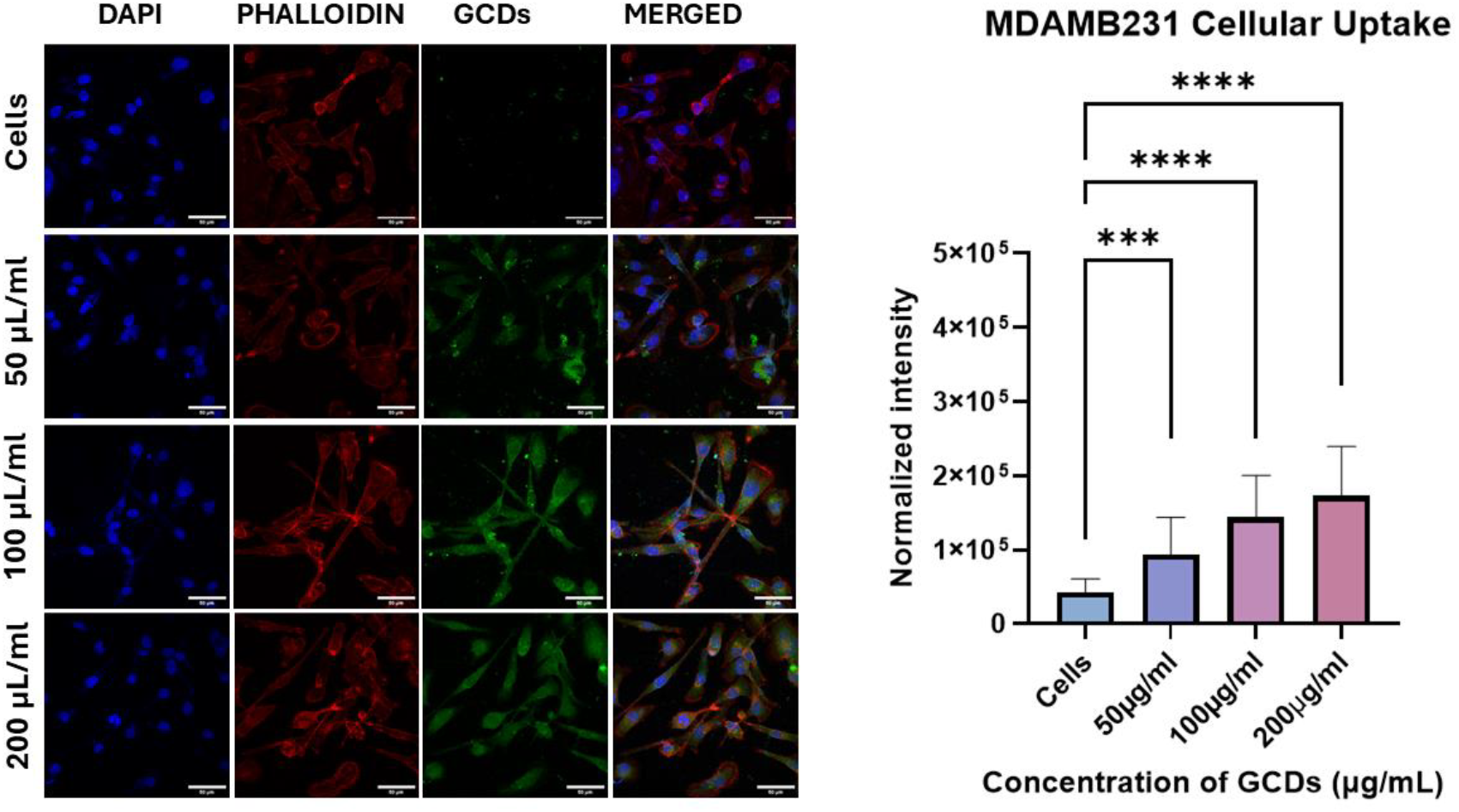
Cellular uptake Of GCDs in RPE1 Cells. The scale bar is 20 µm. uptake concentration of the GCDs 50 μg mL^−1^, 100 μg mL^−1^, and 200 μg mL^−1^ in RPE1 cell. Here, use 2 dyes: DAPI (which stains the nucleus of the cells), and Phalloidin A647 (which stains the cytoskeleton of the cells). Aided to visualize cellular morphology and physiology. The quantification of fluorescent intensity of GCDs in RPE1 cells. Statistical significance was assessed using one-way ANOVA in Prism Software, indicated as **** for p-values < 0.0001, highlighting significant differences among the means (p < 0.05).

**Figure 5.**
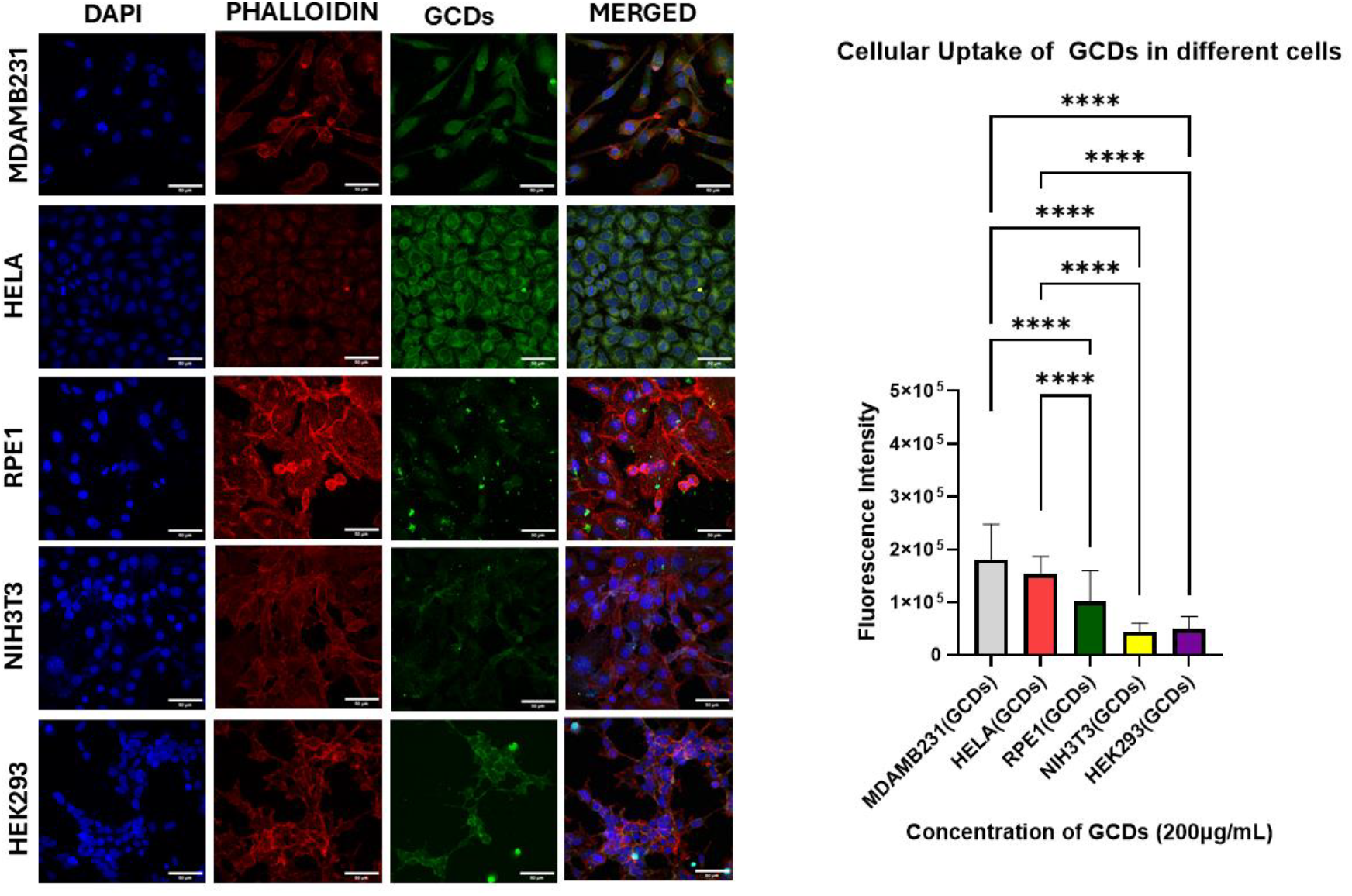
Cellular uptake of GCDs in MDAMB cells.. The scale bar is 20 µm. uptake concentration of the GCDs 50 μg mL^−1^, 100 μg mL^−1^, and 200 μg mL^−1^ in MDAMB231 cell. Here, use 2 dyes: DAPI (which stains the nucleus of the cells), and Phalloidin A647 (which stains the cytoskeleton of the cells). Aided to visualize cellular morphology and physiology. The quantification of fluorescent intensity of GCDs in MDAMB231 cells. Statistical significance was assessed using one-way ANOVA in Prism Software, indicated as **** for p-values < 0.0001, highlighting significant differences among the means (p < 0.05).

MDAMB231 cells. The scale bar represents 50 µm. The quantified fluorescence intensity of the CDs in MDAMB231 was analyzed, and statistical significance was evaluated using one-way ANOVA with Prism software. A p-value less than 0.0001 is indicated by ****, while a significant difference among means is noted at p < 0.05.

In the case of cancerous cells (MDAMB231), there was an increased cellular uptake of GCDs at concentrations of 100 and 200 μg/ml compared to the control and 50 μg/ml. This cellular uptake study is illustrated in the figure above. The cancer cells exhibited higher fluorescence intensity compared to normal cells, likely due to the elevated metabolic rate characteristic of cancerous cells.

### 3.4 Distinguishing between healthy and cancerous cells using the GCDs

GCDs are used as an effective fluorescence probe to distinguish between normal cells and cancerous cells. Five types of cells (including two types of cancerous and three types of noncancerous, for example, MDAMB231, HELA, RPE1, NIH3T3, and Hek293) were separately incubated with 200 µg/ml in a four-well plate, then washed three times with 1X PBS followed by the addition of Triton X-100 and phalloidin A647. Again, it was washed three times in 1X PBS and mounted on the coverslip using DAPI in Mowiol, and the slide was observed in a confocal microscope. As shown in fig., all cancerous cells stained with GCDs emitted bright fluorescence emission, whereas all normal cells emitted weaker fluorescence.

**Figure 6.**Cellular Uptake of GCDs in different types of cells. Distinction of various cancerous cells (MDA-MB-231, HeLa) from normal cells (RPE1, NIH3T3, and HEK293) was achieved using GCDs. The corresponding fluorescence intensities were obtained from confocal microscopy images, with a scale bar of 50 µm. The quantified fluorescence intensity of the carbon dots (CDs) is detailed in MDAMB231. Statistical significance was assessed using one-way ANOVA in Prism Software, with results represented as **** when the p-value is less than 0.0001, indicating a significant difference among means at P < 0.05.

Furthermore, fluorescence signals were quantified using ImageJ software, and the resulting data were plotted for analysis. The fluorescence intensity in cancer cells was found to be significantly higher than in normal cells, which correlated well with the observations from confocal microscopy images. Additionally, the varying fluorescence emission intensities among different cancer cells might be attributed to their distinct ΔΨm membrane potential and metabolic rates. These findings suggest that GCDs hold potential as effective probes for distinguishing between normal and cancerous cells, particularly for early cancer detection.

## 4. CONCLUSIONS AND FUTURE DIRECTIONS

GCDs were successfully synthesized via a reflux method at 130 °C for 12 hours using acetic acid and ascorbic acid as precursors. These GCDs exhibit selective targeting of cancer cells and can differentiate between cancerous and normal cells. The formation of GCDs was initially confirmed through a visual test, where the CDs exhibited green fluorescence under UV light. To further characterize the optical and photoluminescent properties, UV-Vis spectroscopy and fluorescence spectrophotometry were employed. Two prominent absorption peaks were observed at 265 nm and 347 nm, corresponding to electronic transitions. Upon excitation at 347 nm, a strong emission was recorded at 487 nm (∼8036 a.u.), indicating enhanced π→π* transitions or deeper surface state contributions. A secondary green emission peak at 532 nm (∼5912 a.u.) is likely due to n→π* transitions associated with oxygen-containing surface functional groups, such as carbonyls. The consistent emission at 532 nm across different samples confirms that green fluorescence is a characteristic feature of the synthesized GCDs. The size and surface characteristics of the GCDs were analysed using Dynamic Light Scattering (DLS), revealing a hydrodynamic radius ranging from 2–6 nm. Surface charge measurements indicated a pH-dependent trend: as pH increased, the surface charge became more negative. Fourier-transform infrared (FTIR) spectroscopy was used to identify functional groups on the GCD surface, confirming the presence of various oxygen-containing moieties. Atomic Force Microscopy (AFM) further revealed that the GCDs possess a spherical morphology with sizes ranging from 3–10 nm. Biological studies began with cytotoxicity assessments using the MTT assay. In normal RPE1 cells, more than 80% viability was observed at a concentration of 100 µg/mL, whereas in the MDA-MB-231 breast cancer cell line, only 50% viability was observed at 80 µg/mL, demonstrating the GCDs’ anticancer properties. Cellular uptake studies showed that GCDs were internalized more efficiently by cancer cells than normal cells, suggesting selective targeting capabilities. In further experiments involving five different cancerous and normal cell lines incubated with 200 µg/mL of GCDs, cancer cells exhibited significantly higher fluorescence intensity compared to normal cells. This disparity is attributed to the higher membrane potential and metabolic rate of cancer cells, confirming the ability of GCDs to effectively distinguish between cancerous and normal cells.CDs hold great promise as fluorescent probes for biomedical applications, particularly in cancer diagnostics. Future research could focus on enhancing their selectivity and biocompatibility for targeted imaging and therapy. Additionally, CDs offer the potential for exploring the tumour microenvironment, enabling deeper insights into cancer biology and paving the way for early detection and precision treatment.

## Acknowledgements

We sincerely thank all members of the DB Lab at IITGN for their valuable discussions and contributions. We appreciate the Central Instrumentation Facility at IITGN for providing the necessary infrastructure. DB expresses gratitude to ANRF-CRG, GSBTM, and MoES-STARS for their research funding.

## Conflicts of interest

There are no conflicts to declare.

## Notes

### Competing Interest Statement

The authors have declared no competing interest.

